# Improved quantitation in data-independent acquisition proteomics via retention time boundary imputation

**DOI:** 10.1101/2025.05.27.656394

**Authors:** Lincoln Harris, Michael Riffle, Nicholas Shulman, William E. Fondrie, Christine C. Wu, Danielle P. Johnson Erickson, Alec Morimoto, Benjamin A. Shaver, Taylor N. Stein, Ning Cao, Eric C. Ford, William Stafford Noble, Michael J. MacCoss

## Abstract

The traditional approaches to handling missing values in data-independent acquisition (DIA) proteomics are to either remove high-missingness proteins or impute missing values with statistical procedures. Both approaches have their disadvantages: removal can limit statistical power, whereas imputation can introduce spurious correlations or dilute signal. We present an alternative approach based on imputing peptide retention times (RTs) rather than quantitations. For each missing value, we impute the start and end of the peptide’s RT profile (“RT boundaries”), then obtain a quantitative value by integrating the chromatographic signal within the imputed boundaries. We evaluate our method on three distinct datasets and show that RT boundary imputation produces more accurate quantitations than traditional imputation methods and reduces peptide lower limit of quantitation. Additionally, RT boundary imputation allows quantitative ratios to be obtained between experimental groups in cases where this would otherwise not be possible, as demonstrated with matrix matched calibration curve and Alzheimer’s disease datasets. Finally, we show that RT boundary imputation improves the ability to estimate radiation exposure in biological tissues. Our RT boundary imputation method, called Nettle, is available as a standalone tool with an open-source software license.

## 1 Introduction

Advances in mass spectrometry (MS) sample preparation, instrumentation, and data processing have led to significant gains in the number of identifiable proteins in complex mixtures. In spite of this, we are still limited in our ability to accurately quantify proteins. One limiting factor is *missingness*, which refers to proteins that are present in the sample matrix but are not assigned quantitative values. Missingness may be the result of poor peptide ionization, co-eluting peptides, or the inability to confidently assign peptides to spectra [1, 2]. Missingness can limit the statistical power for comparing between samples or experimental groups. Additionally, downstream tasks such as clustering and dimensionality reduction require complete matrices of peptide or protein quantitations.

One approach to handling missingness is to remove high-missingness proteins from downstream analysis. Thresholds for determining whether to exclude a protein are arbitrary, but removing proteins with *>*50% missing values is typical [3, 4]. This practice can remove signal from potentially interesting proteins, especially low-abundance proteins, which are more likely to be missing [5].

Another approach to handling missingness is “plug-in” imputation, in which statistical or machine learning methods are used to estimate missing values based on the observed values alone. Such methods can be thought of as “plug-in” in the sense that imputed values are directly plugged into the matrix of quantitations and used in downstream analysis alongside the observed values. This practice is statistically problematic because there is no notion of the uncertainty associated with imputed values [6, 7]. Plug-in imputation is performed on matrices of quantitations, in which one dimension corresponds to fragments, peptides, or proteins and the other dimension corresponds to MS runs or demultiplexed tandem mass tag (TMT) channels. The canonical example of plug-in imputation is the Perseus strategy of low value imputation [8], in which missing values are replaced by random draws from a Gaussian distribution centered on the lowest observed value in that MS run. MissForest, an approach based on fitting a separate random forest regression model to each missing value [9], is one of the most accurate plug-in imputation approaches [10]. Recently, deep learning (DL) models such as Lupine [11] and PIMMS [12] have emerged, which use deep neural networks (DNNs) to learn patterns of missingness and perform plug-in imputation. While powerful, DL-based methods have longer run times and require specialized hardware (i.e., graphical processing units or GPUs) and technological know-how not available to most labs. Despite its ubiquitousness, plug-in imputation can introduce spurious correlations between proteins and samples [13], as well as noise, further diluting the biological signal [10].

**Table 1.**
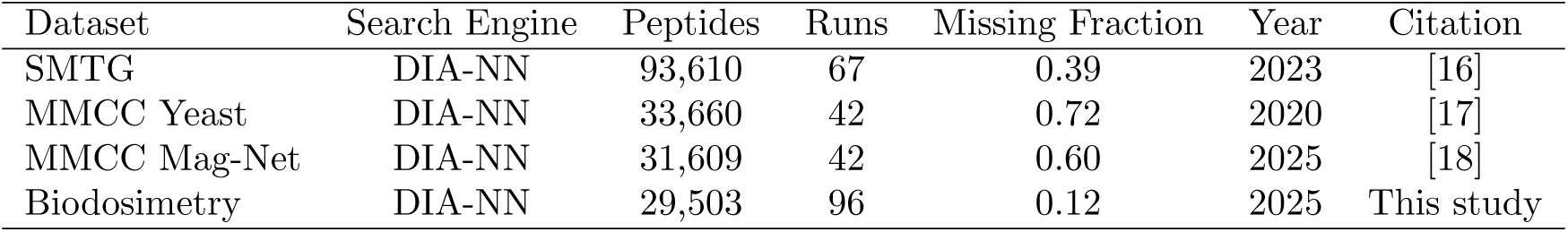
Description of datasets used in this study. SMTG: superior and middle temporal gyri; MMCC: matrix-matched calibration curve.

| Dataset | Search Engine | Peptides | Runs | Missing Fraction | Year | Citation |
| --- | --- | --- | --- | --- | --- | --- |
| SMTG | DIA-NN | 93,610 | 67 | 0.39 | 2023 | [16] |
| MMCC Yeast | DIA-NN | 33,660 | 42 | 0.72 | 2020 | [17] |
| MMCC Mag-Net | DIA-NN | 31,609 | 42 | 0.60 | 2025 | [18] |
| Biodosimetry | DIA-NN | 29,503 | 96 | 0.12 | 2025 | This study |

We address missingness in data-independent acquisition (DIA) proteomics by imputing retention times (RTs) rather than peptide or protein quantitations. Conceptually, our approach borrows RT information from related runs to generate integration boundaries, then performs quantification by integrating the extracted ion current (XIC) within each imputed RT window. We first use a machine learning (ML) approach based on a combination of iterative regularized low-rank matrix completion (soft-impute) [14] and weighted *k*-nearest neighbors [15] to impute missing RT boundaries in a matrix of RT starts and ends. Imputed RT boundaries are then imported into Skyline and XIC-based quantification is performed. The key advantage of our approach is that missing values are replaced with real measured quantities, rather than estimates obtained from statistical or ML procedures.

To demonstrate the utility of our RT boundary impute approach—called Nettle—we apply it to four DIA datasets. These are (i) a dataset obtained from Alzheimer’s disease (AD) clinical samples, derived from the superior and middle temporal gyri (SMTG) brain regions [16], (ii) two matrix-matched calibration curve (MMCC) datasets [17, 18], and (iii) a biodosimetry dataset obtained from mice exposed to varying doses of ionizing radiation. We first show that Nettle produces more accurate quantitations than traditional plug-in impute strategies. We then show that Nettle increases the number of differentially abundant (DA) peptides identified between AD sample groups. These include tryptic peptides from canonical and emerging AD markers such as COL25A1, MAPT, MDK and SMOC1. We show that imputation with Nettle preserves known differences in abundance between AD disease types for two specific Amyloid-*β* tryptic peptides, whereas plug-in impute does not.

In addition, we show that Nettle decreases the lowest sample concentration at which peptides may be reliably quantified, thereby extending the linear dynamic range of peptide quantification. Finally, we show that Nettle improves biodosimetry, or the ability to estimate ionizing radiation exposure in biological tissues [19]. Nettle is an open source implementation of RT boundary imputation and is available on GitHub (https://github.com/Noble-Lab/nettle).

## 2 Results

### 2.1 RT boundary imputation improves quantitative accuracy

To address the accuracy of peptide quantitations obtained with Nettle we first obtained data from Merrihew et al. 2023 [16], a study of clinical AD samples obtained from the superior and middle temporal gyri (SMTG) brain regions. Database search was perfomed with DIA-NN, and an “RT matrix”, in which rows correspond to transitions and columns to RT start or RT end for each MS run, was generated. In the RT matrix, 20% of start-end pairs were masked with either a missing completely at random (MCAR) or missing not at random (MNAR) strategy (Figure 1A). This distinction comes down to whether there is a relationship between missingness and some underlying variable: for MNAR there is, whereas for MCAR there is not [6, 11]. Missingness in proteomics is a combination of MCAR and MNAR, with MNAR typically accounting for the majority [5]. Lower abundance peptides are more likely to be missing.

**Figure 1.**
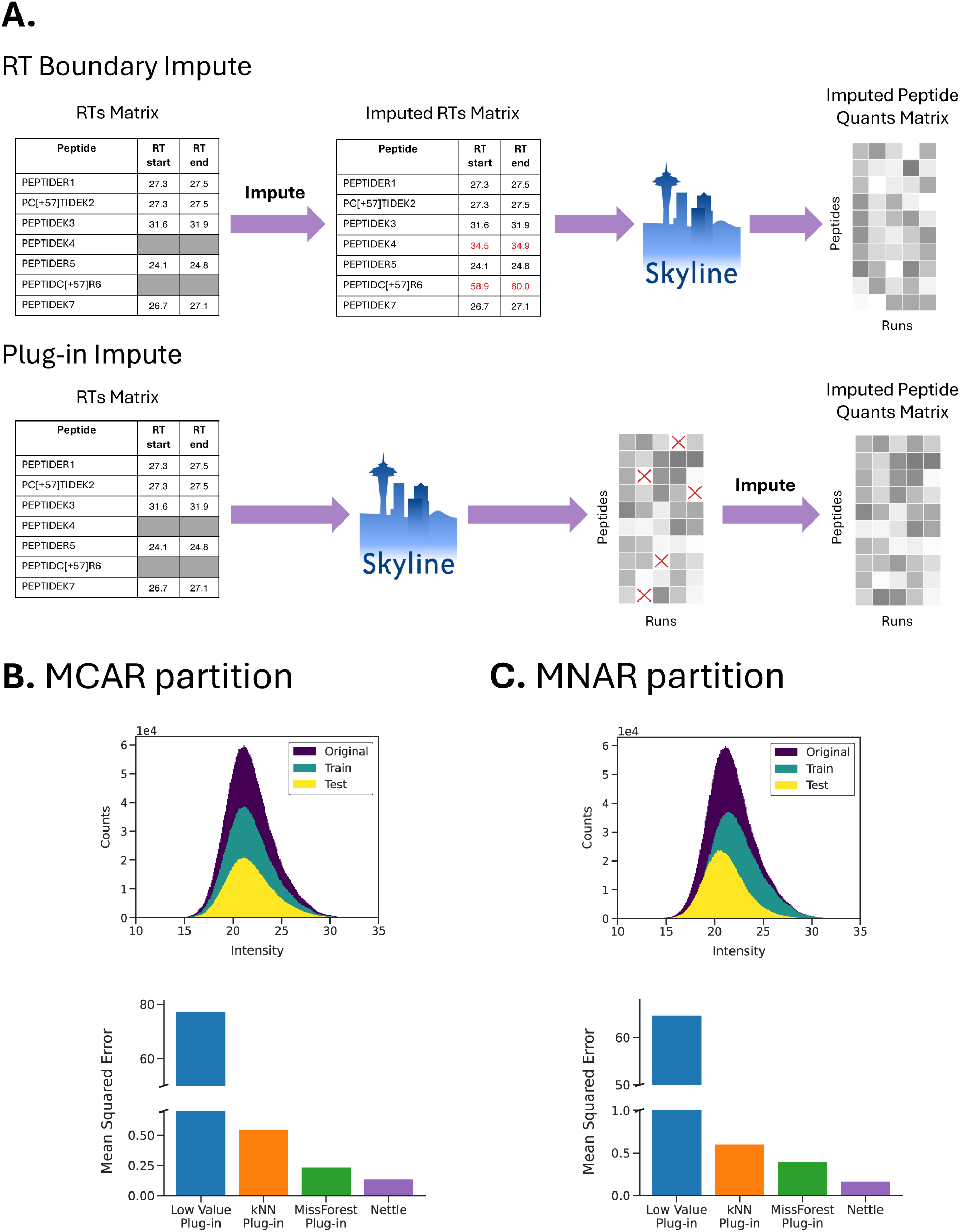
Comparing quantitative accuracy of plug-in to RT boundary imputation with a holdout experiment. **A)** Schematic indicating the difference between RT boundary and plug-in impute. For this experiment, a set of start-end pairs was masked in the RT matrix and imputed with either Nettle or plug-in methods. **B)** For the SMTG dataset, the test set reconstruction accuracy of each method following an MCAR partition. Distributions of the original, train and test sets are shown. **C)** Test set accuracy following an MNAR partition.

We used a previously described MNAR partitioning scheme to simulate a setting in which low-abundance peptides are more likely to be missing, but missingness still occurs across the range of peptide intensities [11]. This entailed generating a quantitative report for an unimputed version of the dataset, then partitioning into train and test such that low-abundance peptides are more likely to end up in the test set. We also performed a standard MCAR partition. The distributions of train and test sets after MCAR and MNAR partitioning are shown in Figure 1B–C. The same mask used to perform this train/test split was then used to hide the corresponding start-end pairs in the RT matrix.

One version of the masked RT matrix was imputed with Nettle, and a second version was left unimputed. For both versions of the RT matrix, Skyline was used to obtain peptide-level quantitative reports. In the plug-in case, this report was then imputed with either KNN, MissForest or low-value impute. KNN [15] is one of the most commonly used plug-in methods [10]; it consists of identifying the *k* most similar MS runs, averaging them and imputing missing values with the mean quantities for each peptide. For each missing value, MissForest [9] fits a random forest regression model based on the observed values, makes a prediction, then moves on to the next missing value. The predicted value is then treated as observed and is included in the training set for the remaining random forest models. “Low value” imputation replaces missing values with random draws from a Gaussian distribution [8].

Figure 1B-C compares the mean squared error (MSE) between observed and imputed quantitations for the test set. Lower MSE indicates more accurate estimates of the held-out peptide quantitations. Because the same set of peptides was held out in both cases, the results are directly comparable between RT boundary and plug-in impute. The results (Figure 1B-C) again indicate that Nettle produces more accurate quantitations than plug-in methods, in both MCAR and MNAR settings. For the MNAR hold-out, Nettle has a test set MSE of 0.17, compared to 0.40 for MissForest and 0.61 for KNN. Again, low value impute performs extremely poorly for both MNAR and MCAR.

We next used an MMCC dataset to assess the quantitative performance of RT boundary impute. An MMCC experiment is a serial dilution series in which human sample is diluted with increasing concentrations of nonhuman background (in this case, yeast) [17]. The background is “matched” in terms of matrix complexity, to prevent undesirable matrix effects during liquid chromatography and electrospray. The advantage of this approach is that the ratios of peptide abundances between samples are known and can serve as ground truth. For example, the ratio for peptide *A* in the 30% dilution sample is given in terms of *A*’s intensity in the 100% (i.e., undiluted) sample: *log*_2_(*A*_30_*/A*_100_).

An MMCC experiment is designed to stretch sample prep methods, instruments and analytical tools to their breaking point. Hence, it is unreasonable to expect any experimental workflow to perfectly capture the expected log ratios, especially for the extreme dilution conditions (e.g., 1%). We focused on two questions with this analysis: (1) Do the boundaries produced by Nettle lead to reasonable peptide quantitations, especially for extreme dilutions, and (2) Does Nettle increase the number of differential peptides that are identifiable between conditions? Figure 2B addresses Question 1: the quantitations obtained with Nettle are close to the expected log fold ratios even for the 3% and 1% dilutions. Supplementary Figure 1 indicates that imputed values are left-skewed relative to the observed, which fits our expectation given that missing values are primarily MNAR, with low abundance quantitations more likely to be missing.

**Figure 2.**
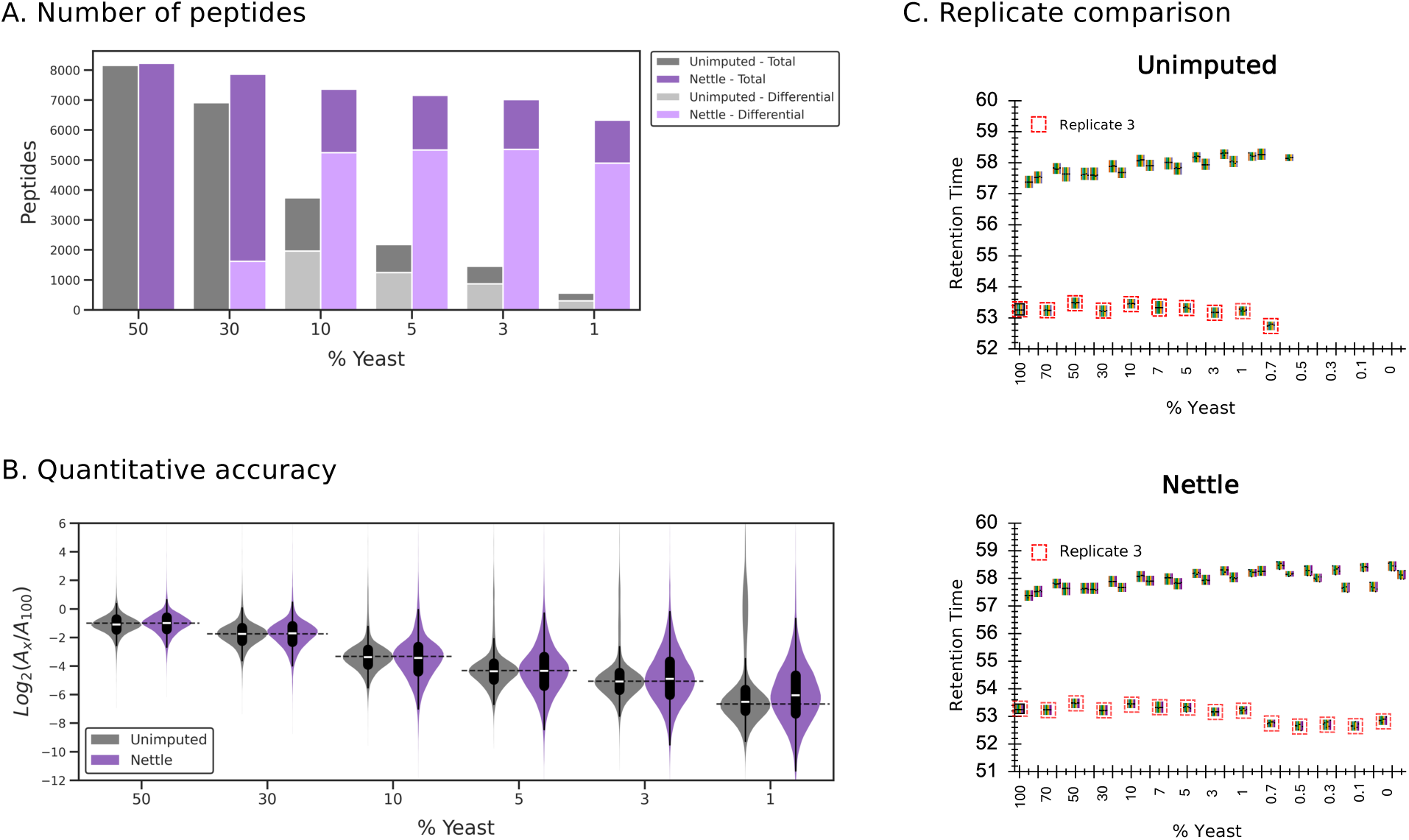
Evaluating quantitative performance of RT boundary impute on a yeast MMCC dataset. **A)** The total number of peptides with quantitations (darker colors) and differential peptides (lighter colors) for each sample in the dilution series, with and without RT boundary imputation. Differential peptides are relative to the 100% (undiluted) condition, for *n* = 3 replicates. In this experimental setup, all peptides are ground truth differential. **B)** Comparing observed and imputed peptide quantitations to the expectation (dashed horizontal lines) for various dilutions. Quantitations are expressed as the log ratio between the value in the dilution and the value in the undiluted (100%) sample (*A_x_/A*_100_). **C)** Retention time comparison across replicates for a single peptide, with and without RT boundary imputation. Three technical replicates were performed; for the third a systemic RT shift was noted (boxed in red). Nettle accounts for this shift, imputing RTs around 53s for the third replicate, and RTs around 58s for the first two.

Figure 2A addresses Question 2, related to quantitative differences between samples. Here we assessed the number of DA peptides identifiable between the 100% samples and each dilution. Two-sample *t* -tests were performed for all peptides; peptides with Benjamani-Hochberg adjusted *p*-values *<*0.05 were considered DA. In the unimputed condition, it is difficult to obtain quantitative differences between peptides because of an abundance of missing values. It is striking that in the unimputed case, there are more DA peptides between 100% and 10% than there are 100% and 1%. This is the opposite of the expectation: greater fold changes should produce more DA peptides. But the excess of missing values makes it difficult to identify DA peptides in the 1% condition. After RT boundary imputation with Nettle, on average 10,105 additional DA peptides are identified per condition, for the six dilutions shown here. For the 30% dilution condition, no DA peptides are detected in the unimputed data, whereas after RT boundary impute 5,649 peptides are detected. It should be stressed that because this is a serial dilution experiment, every DA identification is a true positive.

Another striking observation is that RT boundary imputation appears to implicitly correct for RT shifts. We noticed a significant RT shift in the third replicate of this experiment (shown in Figure 2C, third replicate boxed in red, for tryptic peptide WYPSEDVAAPK). For this peptide, two replicates have RTs around 58s for all samples, while the third elutes near 53s. Nettle accounts for this RT shift, imputing RT boundaries around 58s for the first two replicates and 53s for the third, in the high dilution samples. Note that RT alignment was not performed; Nettle learns the systematic differences between replicates during its training procedure.

We manually inspected some of the Nettle-imputed RT boundaries for the MMCC dataset. We started with the 100% (undiluted) condition, in which the chromatographic peaks should be very well defined, then moved to lower and lower dilutions. Supplementary Figure 2 shows this result for the tryptic peptide LLQFWEPLGET. In the undiluted sample, the peak is well defined and identified by both DIA-NN (left) and DIA-NN + Nettle (right). This is also true in the 30% and 10% conditions. But in the 7% condition, DIA-NN draws a peak boundary that is too narrow, capturing the highest peak but leaving out some of the signal. Nettle appears to capture the entirety of the peak region. In the 5% and 3% conditions, DIA-NN does not assign a peak at all, whereas Nettle does. Thus, Nettle provides a more accurate, empirically observed quantitation for the 7%, 5% and 3% dilutions than library search alone.

### 2.2 RT boundary imputation enables quantitation of additional AD peptides

DA peptides or proteins can indicate biological differences between cells or tissues, serve as biomarkers for disease, or suggest potential drug targets. However, it can be challenging to accurately and reproducibly identify DA peptides between experimental groups, particularly when the number of samples in each group is relatively small. Missingness is part of the reason: if a peptide is completely missing in one group, then it is impossible to calculate a quantitative ratio for it. Put simply, you cannot divide a non-zero quantity by zero. For partially missing peptides, plug-in imputation is often used, which can introduce noise and dilute biological signal [10].

We hypothesized that RT boundary imputation could address this problem. To test this hypothesis, we used the SMTG dataset, which consists of four categories of samples based on behavioral, genetic, and pathological evidence. “ADAD” individuals had causal mutations in *PSEN1*, *PSEN2* or *APP*. This is sometimes referred to as “familial” AD as it can be inherited. “Sporadic” individuals exhibited AD-associated neuropathologic change (e.g., amyloid plaques and TAU tangles) but did not have the canonical *PSEN1*, *PSEN2* or *APP* mutations. “HCF-high” individuals exhibited histopathologic evidence of AD but did not exhibit cognitive symptoms. This is sometimes referred to as “resilient” AD. “HCF-low” individuals exhibited neither histopathologic evidence nor behavioral symptoms.

We identified DA peptides for three comparisons of sample groups: ADAD vs. HCF-low, sporadic vs. HCF-low & ADAD vs. sporadic, with and without RT boundary imputation. In each case, we performed two-sample *t*-tests followed by false discovery rate (FDR) adjustment with Benjamini-Hochberg (BH). Because of the expected biological differences between groups of samples (e.g., it should be relatively easy to identify DA peptides between ADAD and HCF samples [16]), different thresholds were used to define DA peptides. For ADAD vs. HCF-low, we used log_2_ fold change *>*1.0 and FDR-adjusted *p*-value *<*0.01. For the remaining two comparisons, we used log_2_ fold change *>*0.5 and adjusted *p*-value *<*0.05. Because AD is a disease of aberrant proteoforms, we focus exclusively on peptide quantitation—rolling up to the protein level can mask differential signal from important proteoforms such as Amyloid-*β* [20], which gives rise to amyloid plaques and contributes to neuronal death and cognitive decline [21, 22].

Figure 3 shows that for all three comparisons, significantly more peptides with increased abundance were identified after RT boundary imputation. The number of “Nettle unique”, “shared” and “unimputed unique” DA peptides are provided in Table 2. In contrast to ADAD, the genetic underpinnings of sporadic AD are less well understood [23]. Here we attempt to identify DA peptides between sporadic and HCF-low, particularly focusing on the DA peptides that are missed by library search and only identified after post-processing with Nettle (Figure 3C). We identified 40 peptides with increased abundance between sporadic and HCF-low, 30 of which are unique to analysis with Nettle. These unique peptides correspond to AD-associated proteins, including MDK [24], ICAM1 [25] and SMOC1 [26] and NET1 [27]. For the ADAD vs. HCF-low comparison (Figure 3A) we identified expected DA peptides from MDK, APOE, SMOC1 and COL25A1 [16]. The ADAD vs. sporadic comparison (Figure 3B) reveals several potentially interesting APOE DA peptides, although most of these are also found to be DA in the unimputed condition.

**Figure 3.**
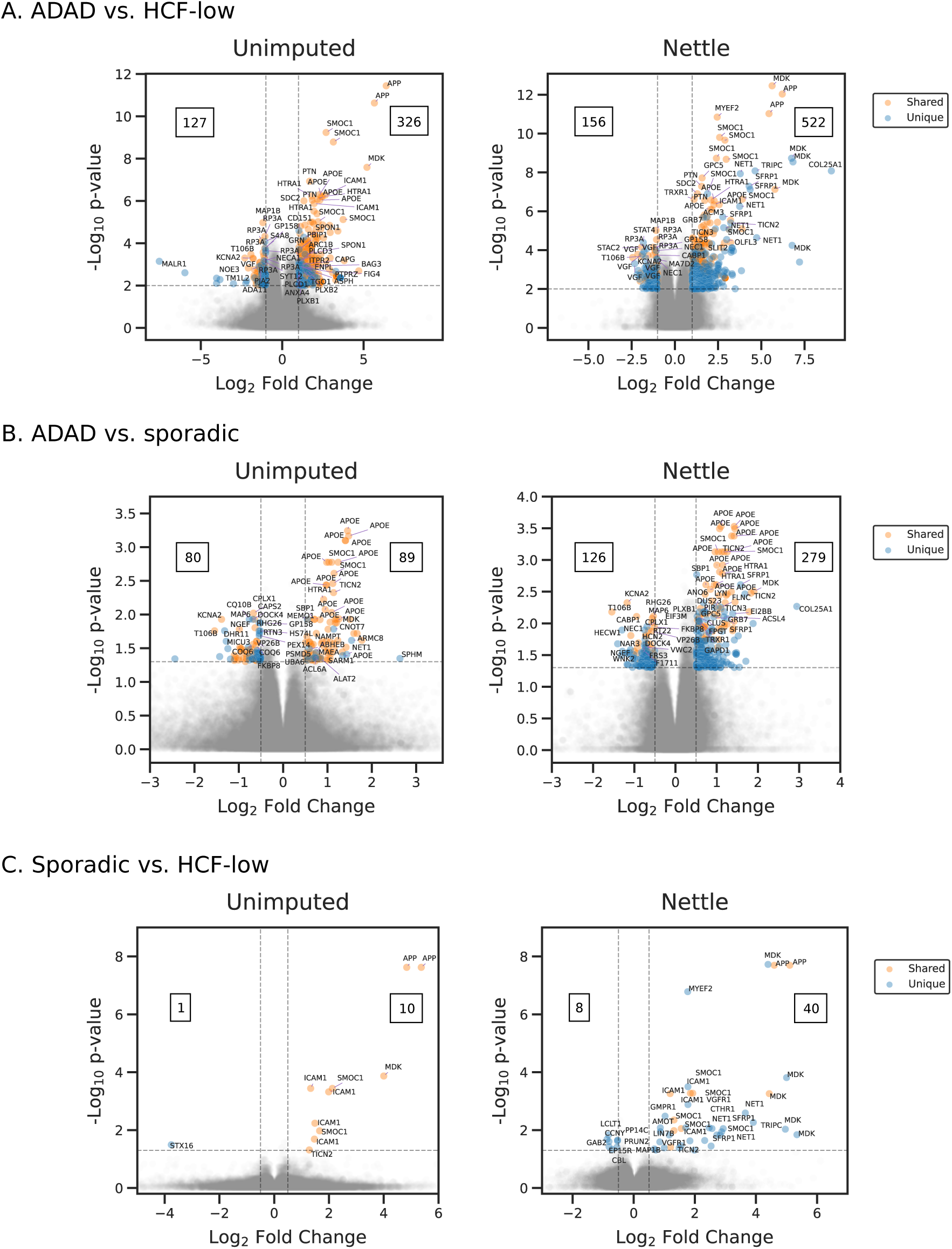
Comparing differentially abundant peptides detected without imputation (left) to RT boundary imputation (right). For a selection of the most significantly differential peptides in the SMTG dataset, the corresponding protein IDs are indicated. Differential peptides unique to either Unimputed or Nettle are shown in blue, shared differential peptides are shown in orange. The total number of peptides with increased or decreased abundances are indicated for each condition.

**Table 2.** Comparing the number of differentially abundant peptides identified with and without RT boundary imputation. ADAD: Autosomal dominant Alzheimer’s disease dementia; HCF: high cognitive function. For the SMTG dataset.

| Comparison | Nettle increased unique | Unimputed increased unique | Increased shared | Nettle decreased unique | Unimputed decreased unique | Decreased shared |
| --- | --- | --- | --- | --- | --- | --- |
| ADAD vs. HCF-low | 270 | 74 | 252 | 90 | 61 | 66 |
| ADAD vs. sporadic | 204 | 14 | 75 | 88 | 42 | 38 |
| Sporadic vs. HCF-low | 30 | 0 | 10 | 8 | 1 | 0 |

To further validate Nettle’s ability to uniquely identify DA peptides, we once again examined individual chromatograms. We focused on two tryptic peptides, INHGFLSADQQLIK and TPSLPTPPTR, of COL25A1 (Collagen-Like Alzheimer Amyloid Plaque) and MAPT (Microtubule-Associated Protein TAU), respectively. COL25A1 is a brain-specific membrane collagen that is implicated in Amyloid-*β* plaque formation [28]. MAPT is a microtubule protein that undergoes complex post-translational modification and splicing and can manifest as aberrant neurofibril tangles, which can contribute to cognitive decline [21]. Supplementary Figure 3A shows that DIA-NN misses a well-defined peak for TPSLPTPPTR (middle panel, sample 16), leading to a missing value for this ADAD sample. Nettle draws reasonable peak boundaries and produces a quantitation for this TAU peptide. INHGFLSADQQLIK is only found to be DA after RT boundary impute with Nettle. As shown in Supplementary Figure 3B, this peak is clearly present in ADAD samples (right) but absent in HCF-low samples (left). DIA-NN does not produce a quantitation for the HCF-low samples; this is a missing value. However, Nettle borrows information from other MS runs to draw reasonable RT boundaries and integrate background signal for the HCF-low samples (Supplementary Figure 3B), assigning a (very low) quantitation to this peptide and allowing us to obtain a quantitative ratio between samples.

Finally, we investigated Amyloid-*β*, one of the most important AD indicators. The parent protein of Amyloid-*β* is Amyloid Precursor Protein (APP), a 750 AA protein which is cleaved by *α*, *β* and *γ* secretases to form the 42 AA Amyloid-*β* peptide [22]. This peptide forms the characteristic dendritic plaques that are associated with AD dementia [29]. Two tryptic peptides of Amyloid-*β*, HDSGYEVHHQK and LVFFAEDVGSNK, have been shown to associate strongly with the familial form of the disease [16]. Merrihew et al. 2023 demonstrated that both tryptic peptides are significantly more abundant in ADAD than sporadic or HCF samples. We wanted to confirm Nettle’s ability to recapitulate this finding in the presence of excess missing values. We used the MCAR procedure to introduce an additional 30% of missing values, then performed imputation with either Nettle or plug-in KNN. MCAR was used to ensure an even distribution of missing values across the two Amyloid-*β* peptides of interest. Figure 4 confirms that after post-processing with Nettle, the SMTG data still capture known AD biology. Significant differences (*t*-test *p*-value *<*0.05) were observed between all groups of samples, with ADAD being the highest, followed by sporadic, then HCF-high. In contract, after plug-in impute with KNN, the expected differences between groups were not found to be significant (Supplementary Figure 4). For HDSGYEVHHQK the difference between ADAD and sporadic is not significant, and for LVFFAEDVGSNK, neither ADAD vs. sporadic nor ADAD vs. HCF-high are significant.

**Figure 4.**
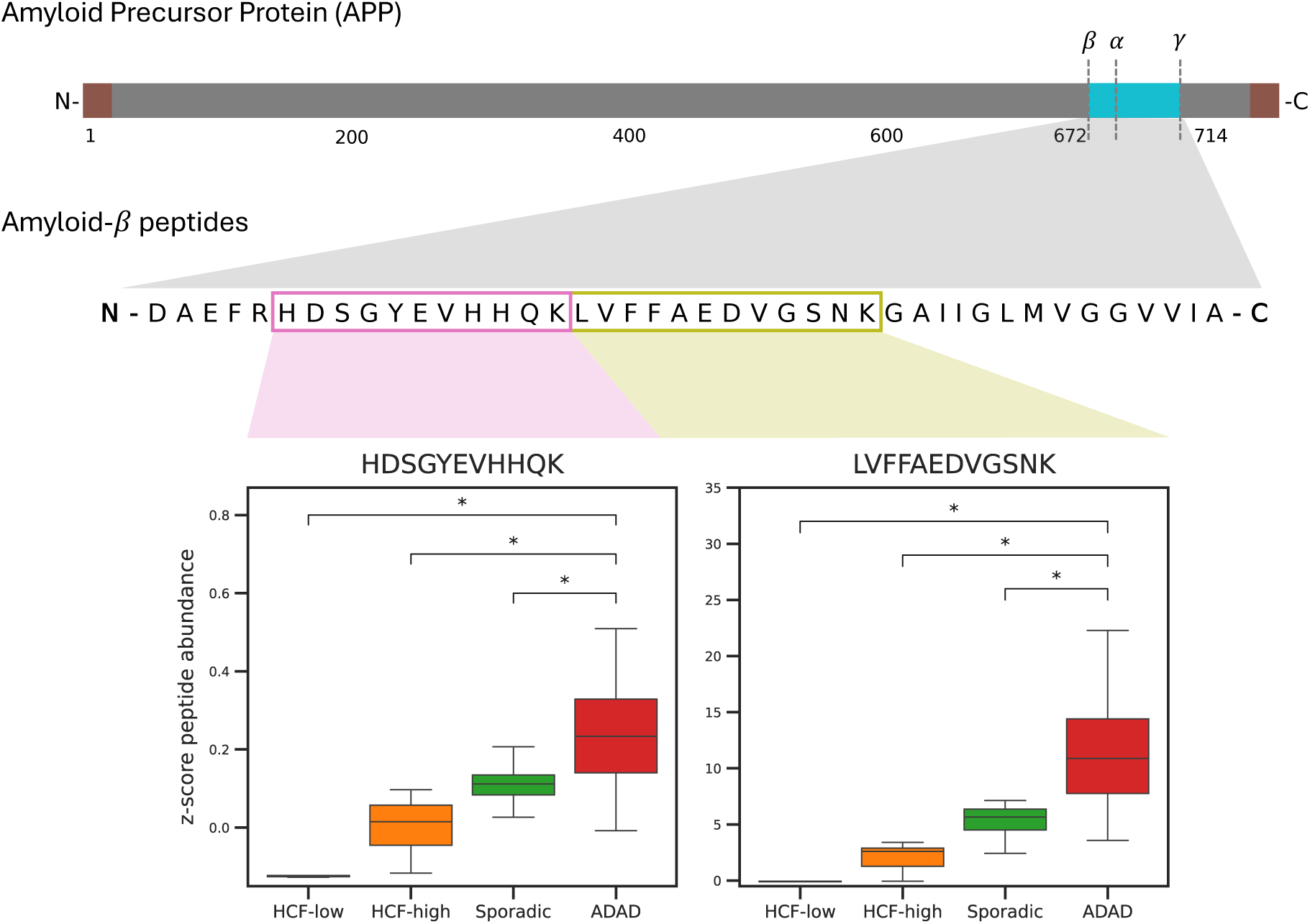
Known indicators of AD pathology are recovered after RT boundary impute. APP is cleaved to produce the 42 AA Amyloid-*β* peptide; tryptic peptides HDSGYEVHHQK and LVFFAEDVGSNK are established indicators of familial AD. 30% additional missing values were introduced, then RT boundaries were imputed with Nettle. *t* -tests were conducted between sample groups, *p*-values *<* 0.05 are starred. For the SMTG dataset.

### 2.3 RT boundary imputation extends the peptide linearly dyanmic range

Many biologically interesting proteins are low abundance, including neurotransmitters in brain, tumor biomarkers in blood, and signaling and transcription factors in virtually all tissues [18]. However, low abundance peptides are more likely to be missing from quantitative proteomics data [5], which can make it challenging to obtain quantitative ratios between groups of samples for these peptides.

Peptides detected via mass spectrometry are only truly quantitative if increases in concentration correspond proportionally to increases in measured signal [17, 30]. Most peptides in an MMCC experiment exhibit a *linear dynamic range* that meets this definition. Above this range, the detector is saturated and increases in concentration do not correspond to proportional increases in signal intensity. The same is true for peptide concentrations below the linear dynamic range: such peptides are below the lower limit of quantification (LLOQ) for that peptide, for that instrument. This phenomenon is sometimes refered to as ratio compression [31]. The LLOQ is distinct from the limit of detection (LOD), which is the concentration of a peptide required for identification [30]. LLOQ is always greater than or equal to LOD.

We used the Mag-Net MMCC dataset to assess whether or not RT boundary imputation can extend the linear dynamic range of peptides. For each peptide, we performed a bi-linear regression designed to model the noise segment—that is, peptide quantities that are below the LLOQ—and linear dynamic range separately. We then calculated the bi-linear turning point, or the analyte concentration at which the noise segment ends and the linear dynamic range segment begins. This idea was introduced by Galitzine et al. 2018 and extended by Pino et al. 2020 [17, 32].

Figure 5 shows bi-linear fits for five peptides for which Nettle extends the linear dynamic range. Quantitations obtained from DIA-NN are shown in blue and quantitations obtained from Nettle are shown in orange. Nettle extends the linear dynamic range past the lowest value obtained with DIA-NN for all five of these peptides. A global version of this analysis is shown in the bottom right of Figure 5. Here the minimum concentration obtained from DIA-NN is shown alongside the Nettle turning point, for all peptides. Generally speaking, Nettle lowers the bi-linear turning point, thereby extending the linear dynamic range and improving our ability to quantify low-abundance peptides.

**Figure 5.**
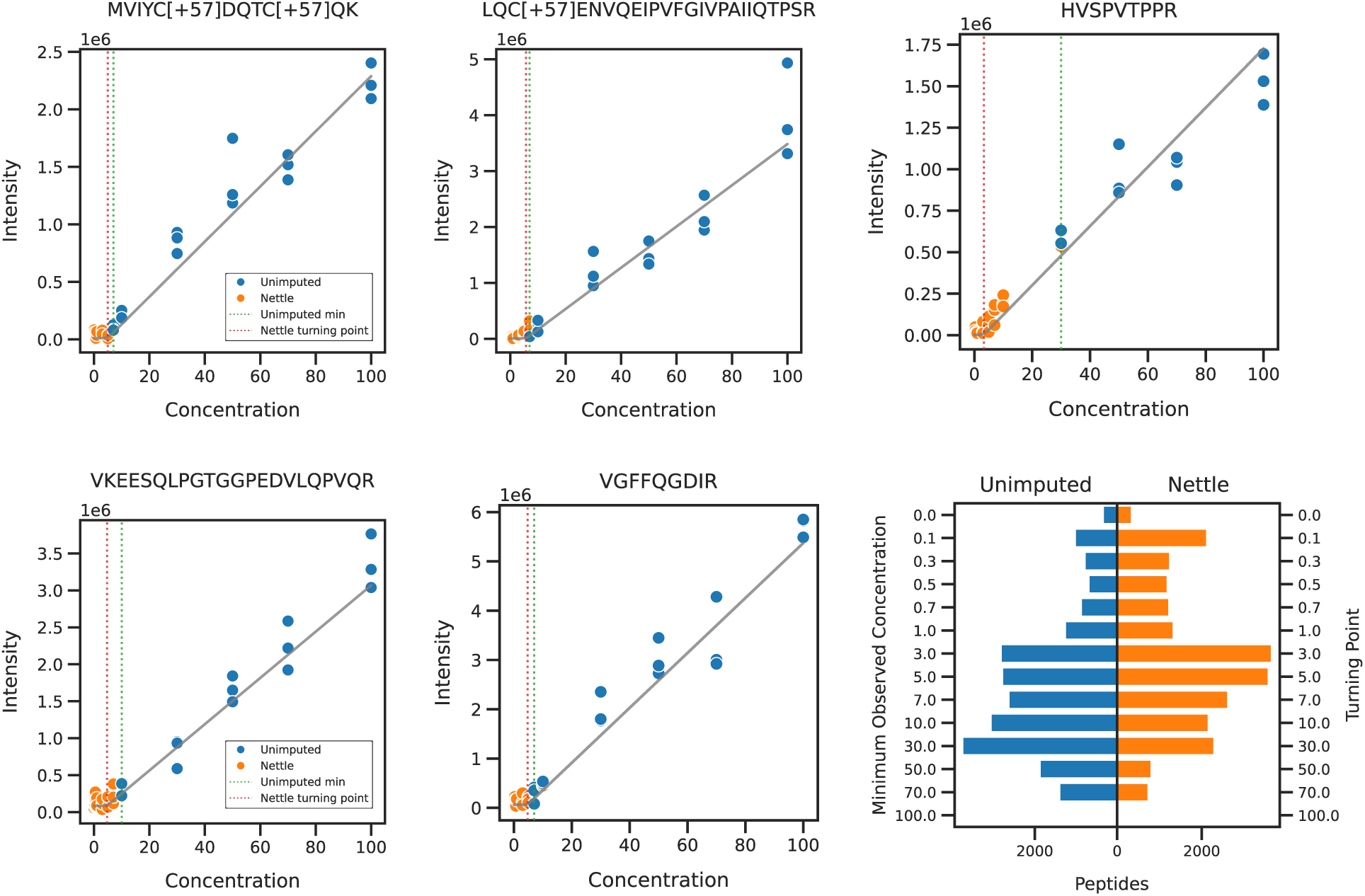
Turning point analysis before and after RT boundary imputation with Nettle. For the Mag-Net MMCC dataset. Nettle imputation was applied and a bi-linear regression model was fit. Calibration curves are shown for five peptides for which Nettle extends the linear range of quantification. Butterfly plot in lower right compares the minimum observed concentration obtained from DIA-NN to the bi-linear turning point obtained with Nettle.

One note is that a significant number of peptides (347 for both unimputed and Nettle) are identified in the 0% dilution. Because no human peptides were added to these samples, all of these identifications are false. In addition to the 1% of false discoveries expected under standard FDR control procedures, many of these are likely to be chicken peptides falsely annotated as human due to high sequence homology. Manual inspection revealed many peptides matching the opposite trend expected in our MMCC experiment—that is, peptides whose concentrations were highest in the 0% dilution condition. These peptides almost certainly belong to chicken.

### 2.4 RT boundary imputation improves biodosimetry

We reasoned that RT boundary imputation might improve the performance of classification or regression models that distinguish between groups of samples. Fitting such models is common practice in proteomics, for example, classifying tumor samples by molecular phenotype to identify potential drug targets [33], identifying coexpression modules in AD samples [27], and associating protein biomarkers with sex in lung disease [34]. The particular use case we chose was biodosimetry, or the ability to estimate ionizing radiation exposure in biological tissues [19].

Data were collected as part of the Intelligence Advanced Research Projects Activity (IARPA) Targeted Evaluation of Ionizing Radiation Exposure (TEI-REX) challenge. The goal of the IARPA TEI-REX challenge was to design an assay for predicting radiation dose from non-invasive samples (e.g., skin disks), particularly for low doses (e.g., *<*1 Gy). The challenge consisted of two phases: in the first, we obtained high-exposure (1–4 Gy) mouse samples and conducted a DIA workflow. We then trained an ensemble of gradient boosted decision trees to predict radiation dose. This analysis allowed us to select a target panel of 92 proteins, corresponding to 1,218 peptides, with high predictive power. In the second phase, we applied the same DIA workflow to a set of low-exposure (*<*1 Gy) samples. We fit an elastic net regularized regression model using the 1,218 peptides in our target panel to predict radiation dose. The median intensity of the target peptides was 10.9, compared to 12.0 for all quantified peptides. The missingness among target peptides was 19%, compared to 12% among all peptides. The results presented here pertain to phase two of the project.

Considering the relatively low intensity and high missingness among target peptides, we reasoned that this analysis would benefit from imputation. We performed plug-in imputation with KNN (*k*=3, i.e., the number of biological replicates for each radiation dose) and RT boundary imputation with Nettle. We performed 10-fold cross validation. The result is shown in Figure 6: the median average error (MAE) between the predicted and true radiation dose decreased from 13.69 for KNN plug-in to 10.02 for Nettle. The Pearson correlation coefficient increased from 0.55 to 0.76. Nettle is better able to quantify the relatively low-abundance peptides in our target panel, leading to improvements in biodosimetry prediction.

**Figure 6.**
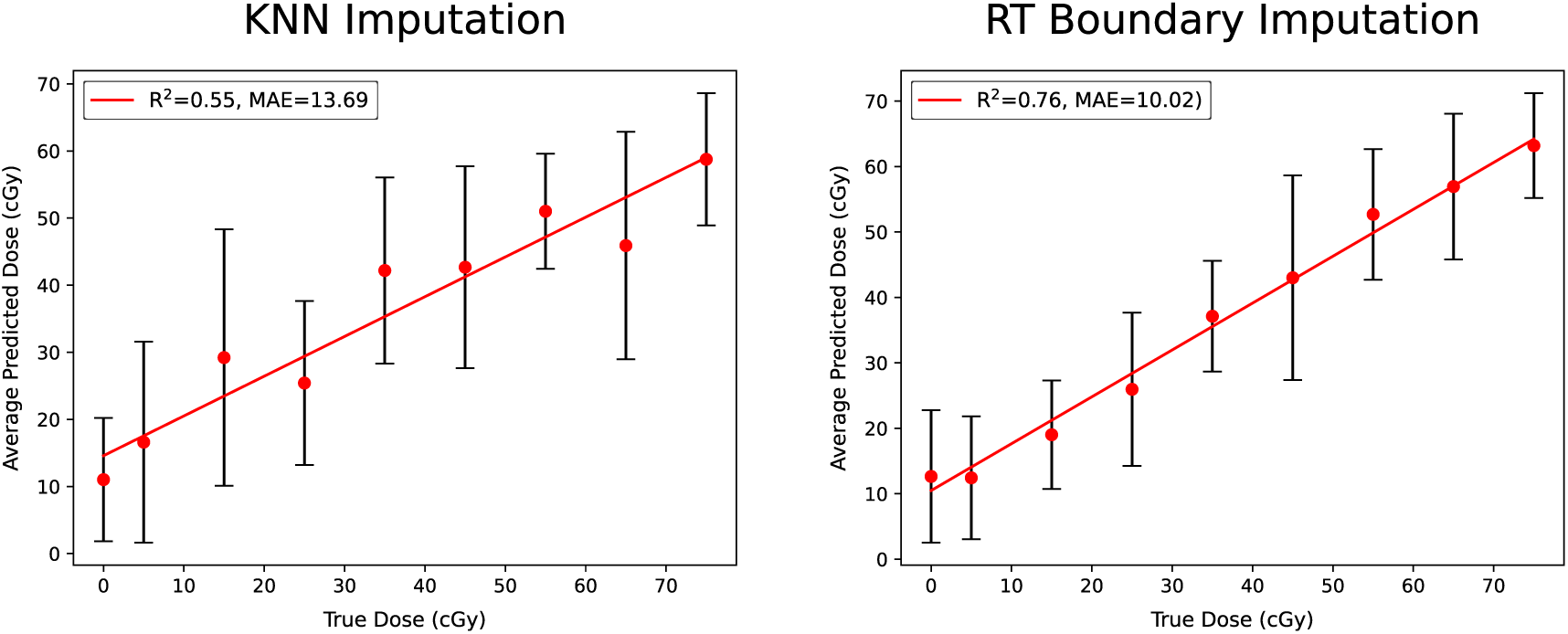
RT boundary imputation improves predictions of radiation dose from DIA proteomic profiles. Mice were exposed to known doses of X-rays (0/sham, 5, 15, 25, 35, 45, 55, 65, and 75 cGy; *≥*8 mice per dose) to generate a standard curve of exposed mouse skin samples. An elastic net regularized regression model was fit to a panel of 1,218 peptides from 92 informative proteins. Regression *R*^2^ and median average error (MAE) are indicated. Best fit lines are shown in red. Error bars indicate one standard deviation from the mean predicted dose. Missing values were handled with either KNN plug-in imputation **(A)** or RT boundary imputation **(B)**.

## 3 Discussion

Missing values limit our ability to draw conclusions from quantitative proteomic datasets. Missing values are traditionally handled with plug-in imputation, which can be statistically problematic because it does not account for the variance in imputed estimates, can wash out biological signal, introduce spurious correlations between samples or proteins, and introduce noise [6, 7, 10]. We present an alternative approach in which missing values are replaced with actual measured quantities derived from the instrument itself. While this procedure can still technically be thought of as imputation, we hope that our reimagined approach to handling missing values can circumvent some of the problems associated with plug-in imputation. We show that our RT boundary imputation method, called Nettle, has practical benefits, including increasing the number of peptides with quantitations, increasing the number of DA peptides, increasing quantitative accuracy and reducing the concentration at which peptides can be reliably quantified. As sample inputs get smaller and experiments get larger, missingness issues will compound [35, 36]. RT boundary impute is a unique and scalable approach to the missingness problem.

One natural question is how RT boundary imputation differs from peptide-identity propagation (PIP), often referred to as match-between-runs (MBR) [37, 38]. PIP arose largely because conventional proteomic workflows are identification-centric: a peptide must first be identified before it can be quantified. PIP addresses missing data by transferring identifications from a “donor” run to an “acceptor” run, matching an unidentified MS1 feature in the acceptor to an identified feature in the donor based on aligned retention time and precursor *m/z*. Critically, this requires a detectable signal in the acceptor run to match to, and FDR is typically controlled only in the donor, not for the transferred identifications. As an important footnote, Solivais et al. 2025 introduce a statistically rigorous method for controlling FDR among transfered identifications [39].

RT boundary imputation operates under a fundamentally different premise than PIP. Rather than attempting to rescue identifications, it provides retention time integration boundaries that enable quantitation of peptides regardless of whether an MS2 signal was detectable. We make no claim that the peptide was identified in the acceptor run — indeed, a key feature of the approach is that it is equally informative when there is no MS2 signal present. In such cases, the resulting near-zero or low-intensity quantitation is itself a meaningful measurement, reflecting that the peptide is absent or below the limit of detection, rather than a missing value to be rescued.

As a way to validate the performance of our method, we examined chromatographs for missing and DA peptides (Supplementary Figure 2, Supplementary Figure 3). We feel that including chromatographic evidence both strengthens our claims and makes our analysis more accessible to mass spectrometry experimenters, who often engage with their data on this level. Furthermore, we encourage users of our tool to examine chromatographic peaks in this manner, for the sake of sanity checking and potentially removing erroneously imputed peaks. The RT boundaries that Nettle draws are not always perfect (Supplementary Figure 3A) and may benefit from manual refinement. We recognize that the gold standard in label-free quantitative proteomics is still manual refinement of peak boundaries. That said, Figure 2D suggest that Nettle implicitly accounts for systematic RT shifts between samples, potentially reducing the amount of manual refinement required.

RT boundary imputation with Nettle identifies more DA peptides between groups of samples than library search with DIA-NN alone. This is especially true for “difficult” comparisons like sporadic vs. HCF-low (Figure 3B). Additionally, we show that for MMCC data, the ability to identify DA peptides between conditions is severely limited by missing values in the high dilution samples. This problem is essentially remedied by RT boundary impute, which identifies on average 10,105 additional DA peptides between conditions (Figure 2C). MMCC serves as a proxy for experiments in which log fold changes between samples are expected but confounded by missingness. We also show that Nettle extends the linear dynamic range of peptides relative to traditional library search alone (Figure 5). The ability to uncover quantitative differences between samples makes Nettle an ideal tool for biomarker discovery. In the future, Nettle may be used to uncover novel biomarkers or potential drug targets in settings where traditional data analysis pipelines fail.

Our biodosimetry analysis (Figure 6) suggests that Nettle can also improve the accuracy of regression or classification models trained on quantitative proteomics data. Such tasks are increasingly important as DIA scales to more and more samples, and may enable classification of tumor subtypes, microbiome samples and single cell proteomes. Especially in contexts where peptides of interest are low-abundance or samples are precious, Nettle may prove a useful tool for data analysis.

Additionally, Nettle increases the number of peptides with quantitations in a mass spectrometry experiment (Figure 2A). While many of these quantities are likely to correspond to integrated background signal, we argue that this is OK: these peptides now have non-missing quantitations and can be included in DA analysis. Without quantitations, these peptides would be either excluded from analysis or imputed with plug-in methods, both of which have well documented issues [10, 13].

Although we focus entirely on DIA-NN in this study, Nettle is also compatible with EncyclopeDIA [40], which generates a spectral library in bibliospec format. Nettle is an open-source, standalone software package available on GitHub, implemented in Python. In the future, Nettle will be directly integrated into Skyline.

## 4 Methods

### 4.1 Data acquisition and processing

We obtained mzML files from three previously published studies: Merrihew et al. 2023 [16], Wu et al. 2024 [18] and Pino et al. 2020 [17]. Here we describe in brief the methods for these three previous studies. We also generated a biodosimetry dataset as part of the Intelligence Advanced Research Projects Activity (IARPA) Targeted Evaluation of Ionizing Radiation Exposure (TEI-REX) challenge. The data generation procedure for the biodosimetry dataset is described below. The number of peptides and runs and pre-imputation missingness fractions for the four datasets are provided in Table 1.

#### 4.1.1 Yeast MMCC dataset

Pino et al. 2020 performed a number of matrix-matched calibration curve (MMCC) experiments to establish analytical figures of merit, including peptide LLOQ, in DIA MS [17]. We analyzed a yeast MMCC dataset from this project. Yeast was diluted with HeLa by volumetric mixing for 14 independent Yeast:HeLa dilution points: (100:0, 70:30, 50:50, 30:70, 10:90, 7:93, 5:95, 3:97, 1:99, 0.7:99.3, 0.5:99.5, 0.3:99.7, 0.1:99.9, 0:100).

Data were acquired with a Thermo Q Exactive HF MS. Thermo raw files from this experiment were obtained from Panorama (https://panoramaweb.org/_webdav/PanoramaPublic/2019/MacCoss-matchedmatrixcalcurves/@files/RawFiles/hela_curves) under ProteomeXchange ID PXD014815 and re-searched with DIA-NN (version 1.8.2).

#### 4.1.2 Mag-Net MMCC dataset

Mag-Net is a procedure for enrichment of extracellular vesicles from plasma using magnetic beads. Wu et al. 2024 used Mag-Net coupled with DIA MS to analyze *>*37k peptides in plasma samples obtained from patients with neurodegenerative disease. We analyzed data from the MMCC experiment that was used as a quality control for the Mag-Net method. Human plasma was diluted with chicken by volumetric mixing for 14 human:chicken dilution points: (100:0, 70:30, 50:50, 30:70, 10:90, 7:93, 5:95, 3:97, 1:99, 0.7:99.3, 0.5:99.5, 0.3:99.7, 0.1:99.9, 0:100). Membrane particles were enriched using the Mag-Net method and performed in triplicate. Samples were analyzed on a Thermo Orbitrap Eclipse mass spectrometer. Thermo raw files were obtained from Panorama (https://panoramaweb.org/_webdav/PanoramaPublic/2023/MacCoss-Mag-NetMethod/@files/RawFiles/MatrixMatchedCalibrationCurve/QuantFiles/) under ProteomeXchange ID PXD042947 [18]. Raw files were converted to mzML with ProteoWizard (version 3.0.23063) [41]. Library search was performed with DIA-NN (version 1.8.2).

#### 4.1.3 SMTG dataset

Merrihew et al. 2023 was a study of individuals enrolled in the Adult Changes in Thought (ACT), the University of Washington AD Research Center (ADRC), or Dominantly Inherited Alzheimer Network (DIAN). Individuals were assigned to four categories based on behavioral, genetic and pathological evidence. “ADAD” individuals had causal mutations in *PSEN1*, *PSEN2* or *APP*. ADAD, or “familial” AD can be inherited and represents only 1% of total cases but can manifest as a debilitating and early-onset form of the disease. “Sporadic” individuals exhibited AD-associated neuropathologic change (e.g., amyloid plaques and TAU tangles) but did not have the canonical *PSEN1*, *PSEN2* or *APP* mutations. “HCF-high” individuals exhibited histopathologic evidence of AD but did not exhibit cognitive symptoms. This is sometimes referred to as “resilient” AD. “HCF-low” individuals exhibited neither histopathologic evidence nor behavioral symptoms. All patients had received a cognitive assessment within two years of death. Samples were obtained from the superior and middle temporal gyri (SMTG) brain regions. In an effort to limit the influence of co-morbidities, rigorous selection criteria were used, resulting in limited sample sizes: 24 ADAD, 18 sporadic, 9 HCF-low and 11 HCF-high, out of a total of 737 pre-screened samples. Tryptic digests from these samples were run on a Thermo Orbitrap Fusion Lumos mass spectrometer. The Thermo raw files were obtained from Panorama (https://panoramaweb.org/ADBrainCleanDiagDIA.url) and converted to mzML with ProteoWizard (version 3.0.23063)[41].

#### 4.1.4 Biodosimetry dataset

Male C57BL/6 11-week-old mice were sourced from The Jackson Laboratory and housed under standard conditions with food and water *ad libitum*. Animals received a single total body dose of 5, 15, 25, 35, 45, 55, 65, or 75 cGy using a Small Animal Radiation Research Platform (SARRP) Xray source (Xstrahl, Inc). Dose rates were calibrated based upon the procedures described in American Association of Physicist in Medicine (AAPM) Task Group Report 61 (TG-61) (Ma et al. 2001) with regards to the following conditions: X-ray tube potential was 220 kV and the half value layer (HVL) was 0.67 mm copper (Cu) with a nominal field size of 13x17 cm at the treatment plane [42]. Doses were measured at 0.5 cm depth in solid water phantom using a Preston-Tonks-Wallace (PTW) model N30013 Farmer-type ionization chamber and Standard Imaging, Inc, model SUPERMAX electrometer. The chamber and electrometer underwent calibration at the University of Wisconsin Medical Radiation Research Center Accredited Dosimetry Calibration Laboratory (UWMRRC ADCL) using a beam quality of UW200-M and UW150-M. Two different dose rates were used: 300 cGy/min (tube current 10.7 mA, source-to-surface distance 33 cm) and 30 cGy/min (4.6 mA, SSD 76 cm). Control animals were sham irradiated under anesthesia. Animals were euthanized by cervical dislocation under isoflurane anesthesia 6 days post-irradiation. A minimum of *n* = 8 mice were collected per exposure. Immediately following euthanasia, mice were shaved and pelts were dissected using a dorsal midline incision. 5-mm skin punches were collected from the dissected pelts using disposable biopsy punches (Integra Miltex, catalog #33-35), transferred into sterile 1.5 mL microcentrifuge tubes, frozen on dry ice, and stored at -80°C.

Pelt punches were probe sonicated in lysis buffer (100 mM Tris, pH 8.5/2% SDS + HALT Protease and Phosphatase Inhibitor Single-Use Cocktail) at room temperature (Fisher Scientific Model 100 Ultrasonic Dismembrator with Microtip Probe). The resulting supernatants were solubilized and reduced in 50 mM Tris, pH 8.5/1% SDS/10 mM Tris (2-carboxyethyl) phosphine (TCEP) with 800 ng enolase standard added as a process control as previously described in Tsantilas et al. 2024 [43]. Samples were alkylated with 15 mM iodoaceamide in the dark for 30 minutes and then quenched with 10 mM DTT for 15 minutes. The samples were processed using protein aggregation capture (PAC) on magnetic microparticles as previously described in Batth et al. 2019 with minor modifications [44]. Briefly, the samples were adjusted to 70% acetonitrile, mixed, and then incubated for 10 minutes at room temperature to precipitate proteins onto the bead surface. The beads were washed 3 times in 95% acetonitrile and 2 times in 70% ethanol for 2.5 minutes each on magnet. Samples were digested for 1 hour at 47°C in 50 mM Tris, pH 8.5 with Thermo Scientific™ Pierce™ porcine trypsin at a ratio of 1:20 trypsin to protein. The digestion was quenched to 0.5% trifluoracetic acid and spiked with Pierce Retention Time Calibrant (PRTC) peptide cocktail (Thermo Fisher Scientific) to a final concentration of 50 fmol/*µ*L as a process control as previously described in Tsantilas et al. 2024 [43]. Peptide digests were then stored at -80°C until LC-MS/MS analysis.

### 4.2 Liquid chromatography-mass spectrometry for the biodosimetry dataset

Peptide digests from the biodosimetry dataset were separated using a Thermo Scientific Vanquish Neo UHPLC system. Peptides were separated using a 21-minute analytical gradient at a flow rate of 1.0 *µ*L/min. Mobile phase A was 0.1% formic acid in water and mobile phase B was 0.1% formic acid in acetonitrile. Following a loading step at 1.3 *µ*L/min and 3% B, the gradient ramped from 4.5% B to 38% B over 21 minutes. A column wash was then performed by increasing to 50% B at 22.5 minutes and then to 99% B at 23 minutes, held for 1 minute, all at 1.3 *µ*L/min. The total run time was 24 minutes.

Data were acquired on a Thermo Scientific Orbitrap Astral mass spectrometer operating in positive ion mode using data-independent acquisition (DIA) as described previously in Heil et al. 2023 [45]. MS1 survey spectra were acquired in the Orbitrap at 240,000 resolving power over a mass range of 375–985 *m/z* using the “Standard” AGC target. DIA scans were collected using 4 *m/z* -wide isolation windows spanning a precursor mass range of 400–900 *m/z* with 124 scan events per cycle. Window placement optimization was enabled [46]. Fragment ions were generated by beam-like collision induced dissociation (HCD) at a normalized collision energy of 27% and using a default charge-state of 2. MS2 spectra were acquired in the Astral analyzer over a scan range of 200–1960 *m/z* with centroid data type. The maximum injection time was set to 10 ms, the normalized AGC target was 250%, and the RF lens was set to 50%.

### 4.3 RT boundary imputation

Each mzML file was searched with DIA-NN v1.8.2 [47] using the following settings: *unimod4* ; *qvalue* 0.01; *cut K*, R*, !*P* ; *reanalyze*; *smart-profiling*. Spectral libraries in bibliospec format, generated with DIA-NN, were converted to CSVs in which rows correspond to transitions and columns correspond to RT start or RT end for each MS run (referred to as the “RT matrix”). The RT matrix was winsorized to limit the influence of extreme values, which presumably correspond to contaminants. The 10% most extreme values were eliminated. Following winsorization, an additional filtering step was performed based on the number of observed values in the RT matrix. Peptides with fewer than five non-NaN start-end pairs were discarded. The goal here was to avoid imputing RTs for peptides that are almost never observed and may not have reliable RTs for the model to learn.

We developed Nettle, a method for RT boundary imputation. Nettle ensembles two complementary imputation approaches and operates solely on the RT matrix. The first approach is distance-weighted *k*-nearest neighbors (wKNN), for which the peptide RTs were features and runs were samples, and the distance metric was Euclidean. Scikit-learn’s (v1.5.2) KNNImputer implementation was used. For the majority of analyses, *k* was set to eight. For experiments with replicate structure, we generally recommend that *k* be set to the number of technical or biological replicates. We ran KNNImputer separately for RT starts and RT ends. If, for a given peptide, the imputed RT start is greater than or equal to the RT end, the values were swapped.

The second method was soft-impute [14], a fast-converging, low-rank matrix completion algorithm that iteratively replaces missing values with those obtained from a nuclear norm-regularized low-rank singular-value decomposition (SVD) dense matrix. A custom python implementation of soft-impute was used, available as part of the Nettle python package (https://github.com/Noble-Lab/nettle). The default hyperparameters were used (*shrinkage value:* None, *max rank:* None, *max iters:* 100, *tolerance:* 1e-4). The relative contributions of wKNN and soft-impute is a user-tunable parameter with a default of 0.5, indicating equal contributions. Similar to the wKNN approach, if a peptide’s RT start is greater than or equal to its corresponding RT end, then both are set to NaN.

After imputation, the completed RT matrix was written back to a bibliospec spectral library, then read into a Skyline document along with the original mzML files and FASTA file. For the SMTG and biodosimetry datasets, the following Skyline peptide settings were used: *Normalization Method:* Equalize Medians; *MS level:* 2. For the MMCC dataset, the following Skyline settings were used: *Regression Fit:* Bilinear; *Normalization Method:* None; *Regression Weighting:* None; *MS level:* 2; *Calculated LOD by:* Bilinear turning point. The following transition settings were used: *Library:* 6 product ions, 4 minimum product ions; *Full-Scan: Retention Time Filtering:* Use only scans within 3 minutes of MS/MS IDs.

We exported Skyline reports of peptide abundances, where “abundance” refers to the sum of peak areas for all transitions of a given peptide after background subtraction. RT alignment was not performed. We reasoned that a clustering algorithm such as distance-weighted KNN should be able to account for systematic differences in RTs between samples without explicit RT alignment. The Skyline documents used in this study can be found on Panorama Public (https://panoramaweb.org/GS4ghw.url) and ProteomeXchange (PXD076519) [48]. The code to reproduce the figures can be found at: https://github.com/Noble-Lab/rt-boundary-manuscript.git.

For SMTG, peptide quants were median normalized, then batch corrected with ComBat [49]. For the biodosimetry dataset, median absolute deviation (MAD) normalization was performed with the Pronomos Python package.

### 4.4 Turning point analysis

For each peptide, we fit a piecewise function with slope = 0 below a specified turning point value, and slope *>* 0 above the turning point. Reasonable initial estimates for parameters were obtained, then non-linear least squares regression was used to fit the piecewise function. The regression model was weighted by uncertainty estimates for each point.

After curve fitting, estimated turning points were replaced with the next largest concentration in the MMCC experiment. If an estimated turning point was greater than the lowest observed DIA-NN quantification for that peptide, the turning point was replaced with the latter value. Peptides were then ranked according to the slope of the linear dynamic region obtained without Nettle imputation. In an effort to remove interfering chicken peptides, the bottom 10% of peptides ranked by slope were excluded from analysis.

## Acknowledgements

The authors thank Bo Wen and Chris Hsu of the University of Washington’s Department of Genome Sciences for helpful discussions. This work was supported in part by National Institutes of Health grants F31AG082395, U19AG065156, and R24GM141156. Support was also provided by the Intelligence Advanced Research Projects Activity (IARPA) TEI-REX program through the Army Research Office contract W911NF2220059. The views and conclusions contained should not be interpreted as necessarily representing the official policies, either expressed or implied, of ODNI, IARPA, ARO, or the U.S. Government. The U.S. Government is authorized to reproduce and distribute preprints for governmental purposes notwithstanding any copyright annotation therein. This work is supported in part by the University of Washington’s Proteomics Resource (UWPR95794).

## Author Information

### Notes

The MacCoss Lab at the University of Washington has a sponsored research agreement with Thermo Fisher Scientific, the manufacturer of the mass spectrometry instrumentation used in this research. Additionally, Michael MacCoss is a paid consultant for Thermo Fisher Scientific.

---

**Supplementary Figure 1.**
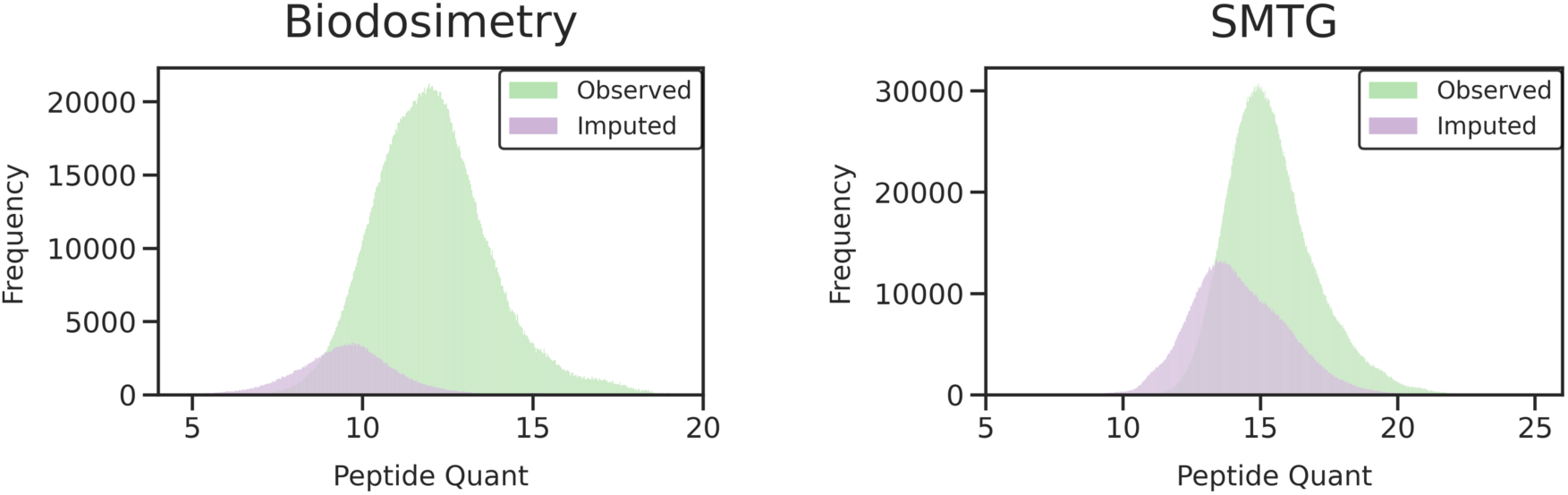
Distribution of observed and imputed quantitations for the biodosimetry and SMTG datasets. Quantities were imputed with Nettle. Peptide intensities have been *log*2 transformed.

**Supplementary Figure 2.**
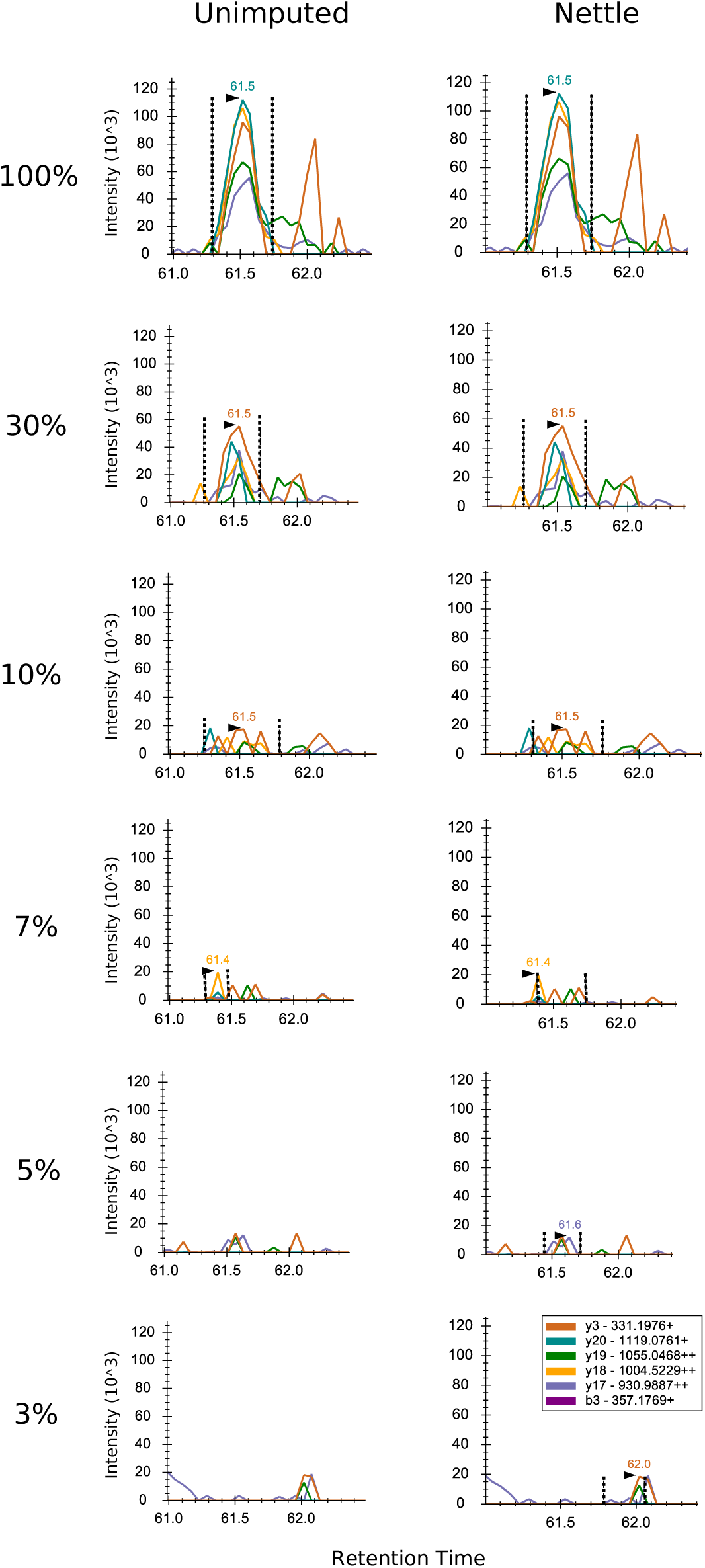
Chromatograms for the peptide LLQFWEPLGETR at different dilutions in the Mag-Net MMCC experiment. Library search was performed with DIA-NN; chromatograms were visualized with Skyline. Right: missing RT boundaries were imputed with Nettle. Percentages on the left indicate the fraction of human peptides relative to matrix-matched background in the sample. Vertical bars indicate RT integration boundaries for that peptide.

**Supplementary Figure 3.**
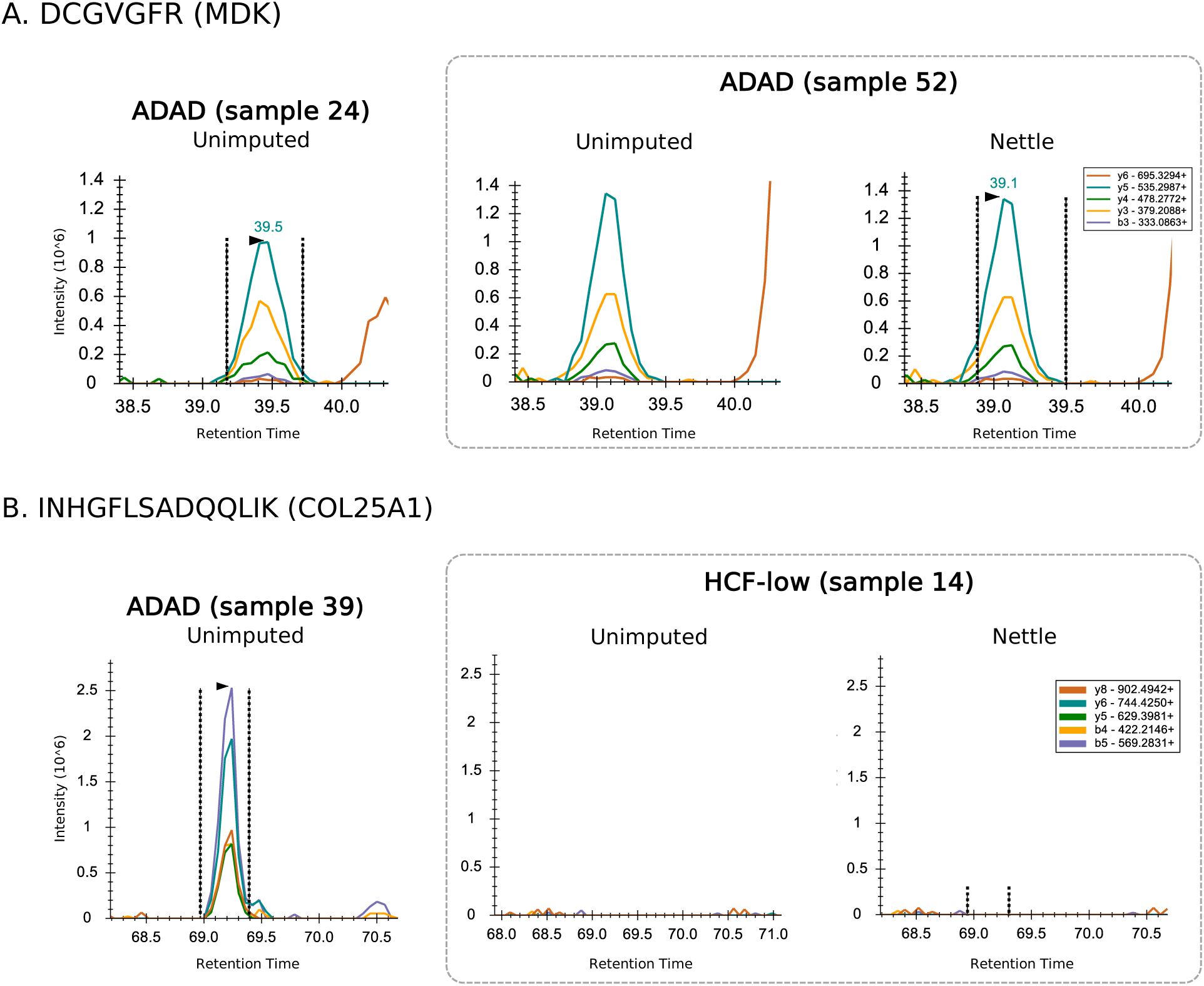
RT boundary impute enables quantitation of Alzheimer’s disease peptides. **A)** Chromatographic peaks for a tryptic peptide of COL25A1. All three panels from ADAD samples. Vertical bars indicate RT integration boundaries for that peptide. The peak in sample 52 is missed by DIA-NN, but imputed with Nettle. **B)** Chromatograms for a tryptic peptide of TAU. Left and center panels are from HCF-low samples, right is from ADAD. Sample 14 was a missing value with DIA-NN, but is assigned a non-zero quantitative value with Nettle.

**Supplementary Figure 4.**
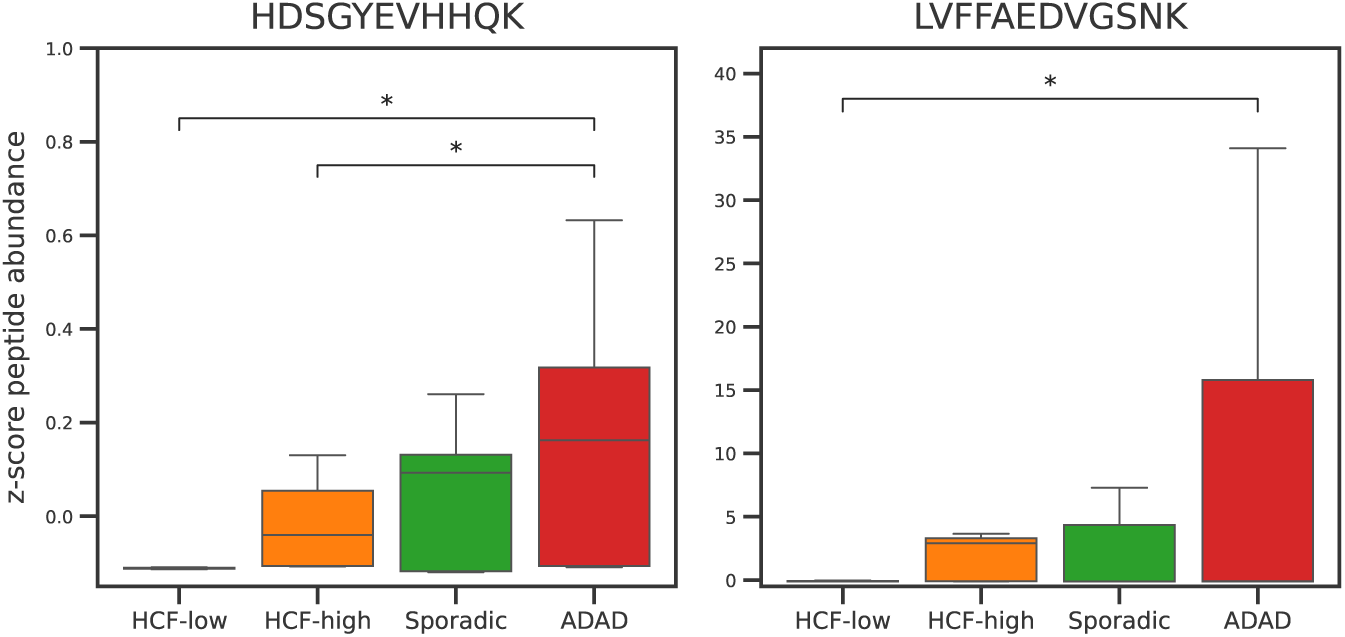
KNN plug-in imputed does not differentiate sample groups for known indicators of AD pathology. The tryptic peptides HDSGYEVHHQK and LVFFAEDVGSNK are known indicators of ADAD. 30% additional missing values were introduced and quantitations were imputed with plug-in KNN. *t* -tests were conducted between groups of samples, *p*-values *<* 0.05 are starred. For the SMTG dataset.

